# Using AlphaFold Multimer to discover interkingdom protein-protein interactions

**DOI:** 10.1101/2024.06.14.599045

**Authors:** Felix Homma, Joy Lyu, Renier A. L. van der Hoorn

## Abstract

Structural prediction by artificial intelligence (AI) can be powerful new instruments to discover novel protein-protein interactions, but the community still grapples with the implementation, opportunities and limitations. Here, we discuss and re-analyse our in-silico screen for novel pathogen-secreted inhibitors of immune hydrolases to illustrate the power and limitations of structural predictions. We discuss strategies of curating sequences, including controls, and reusing sequence alignments and highlight important limitations originating from platforms, sequence depth and computing times. We hope these experiences will support similar interactomic screens by the research community.

---

Artificial intelligence (AI) is making a deep impact in molecular sciences, including plant science. The prediction of protein structures with Alphafold2 has caused a paradigm shift, but many scientists are still grappling with its implementation, opportunities and limitations. We recently demonstrated the use of AlphaFold Multimer (AFM, Evans et al., 2022) in identifying novel interactions between plant proteins and proteins secreted by microbial pathogens (Homma et al., 2023). We screened 11,274 protein pairs with AFM for possible complexes between 1,879 small secreted proteins (SSPs) of pathogens and six secreted immune hydrolases of tomato. Biochemical assays were used to confirm that four of these SSPs are indeed novel inhibitors of secreted immune protease P69B of tomato (Homma et al., 2023). Whilst we are validating the other predictions of this screen and conducting similar screens to explore other pathosystems, we experienced high public interest in the procedures that we followed. Here, we would like to share our experiences and add additional data interpretations to support the development of similar *in-silico* interactomic screens in the research community. The discussed choices and parameters in input, platform and output are summarised in **Figure 1**.

**Figure 1.**
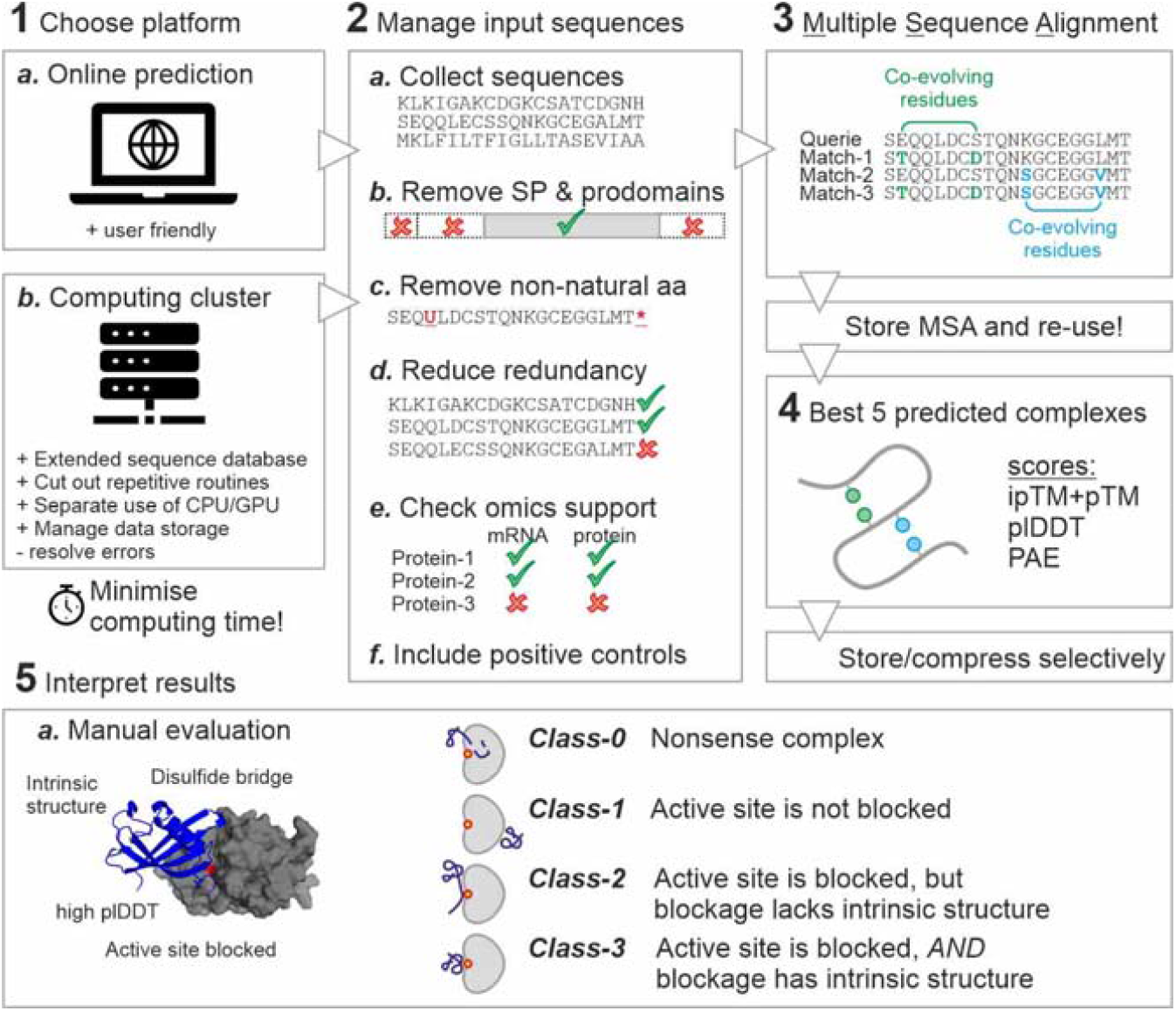
Flow chart of AlphaFold Multimer. These topics are discussed in the main text.

## Start with ColabFold online

Researchers that have never used AFM before can start with the online implementation of ColabFold (Mirdita et al., 2022), which is a free web-based service that can be accessed via ‘structure prediction’ in the ‘tools’ tab in ChimeraX (Pettersen et al., 2021) or as a Jupyter Notebook in Google Colaboratory (Mirdita et al., 2022). ColabFold predicts structures on Google servers and a link to your Google account will be used to grant access to a reasonable amount of trial time, enough to model e.g. 20 complexes. After that, the application slows down and this barrier can be overcome by buying credits that will not only extend your access, but will also give you access to faster processing units. ColabFold online is great for many proteins but we noticed important limitations in throughput and the quality of complex predictions (see below).

### Use a computing cluster for screens

We perform(ed) the larger AFM screens on a computing cluster of the University of Oxford (ARC, Richards, 2015). The modularity of AFM on a computing cluster allows the allocation of different tasks to different processing units and to feed AFM with different input sequence databases. Running AFM on the cluster, however, requires extensive experiences in Linux environments and a strong resilience to resolve a diverse range of error messages. Computing clusters enable high throughput predictions of protein structures, depending on availability of the required hardware, especially the access to Graphics Processing Units (GPUs). Good access to computing clusters (e.g. access to over 70 GPUs in parallel) will enable to predict ∼1,000 protein pairs per day but computing clusters can be expensive to use and/or can have long waiting queues due to increasing popularity of machine learning tools in biological research. We noticed striking differences in ipTM+pTM and average plDDT scores between the online AFM implementation via ColabFold and AFM on the cluster, illustrated with predictions of known SSP-protease complexes and their negative controls and our four validated novel SSP-P69B interactions (Homma et al., 2023) (**Figure 2**). Importantly, three of the four novel P69B inhibitors would not have been identified using the free online implementation of ColabFold because their highest ipTM+pTM scores are below 0.75 (**Figure 2**). We suspect this is mostly due to the size of the underlying database, see below (*Try to get MSA >100*).

**Figure 2.**
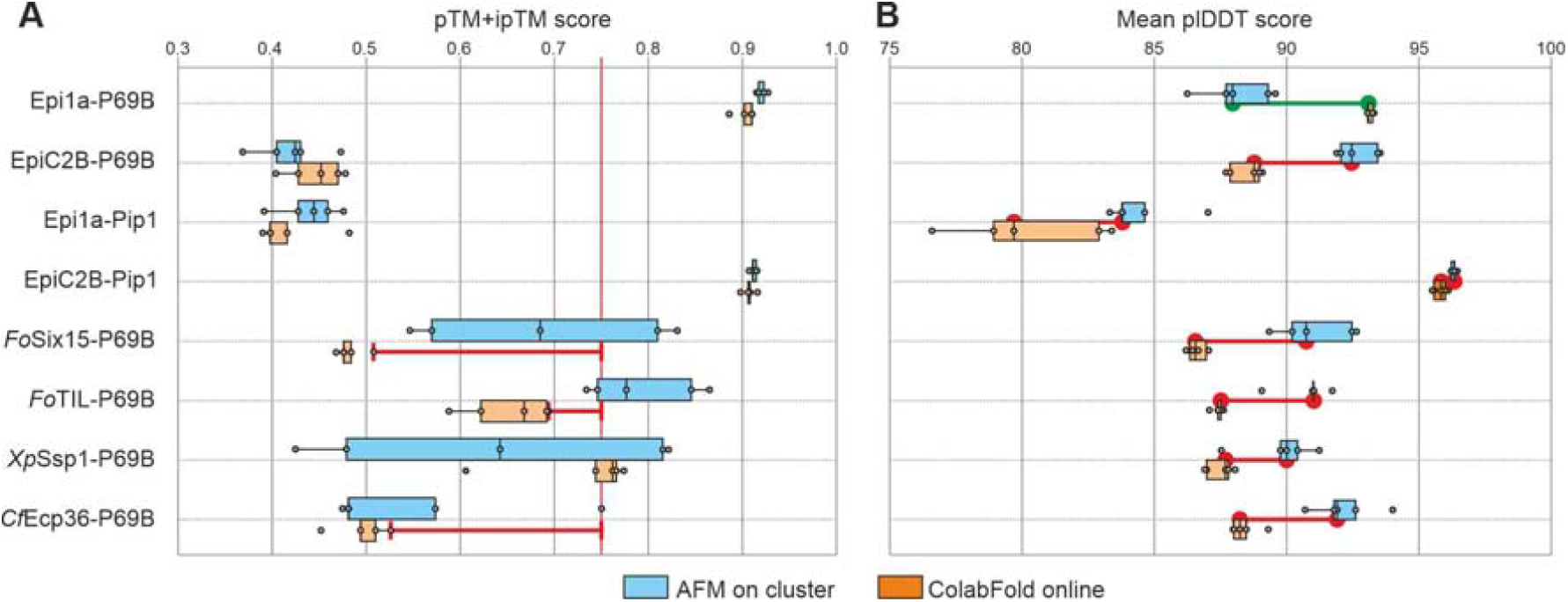
Lower hit rates for free online implementation of ColabFold in SSP-hydrolase screen. Structural models of known and novel SSP-hydrolase pairs and negative controls (Epi1a-Pip1 and EpiC2B-P69B) were generated with the online implementation of ColabFold and using the ARC high-performance computing cluster, plotting their ipTM+pTM scores **(A)** and plDDT scores **(B)**. Boxplots summarise the distribution of complex scores, with the edges showing upper and lower quartile separated by the line representing median values. Note that the highest-scoring model for three of the four novel P69B inhibitors (*Fo*Six15, *Fo*TIL and *Pi*Ecp36) would not score above the 0.75 threshold when predicted online via ColabFold.

### Small sequences model faster

Before setting up screens, it is important to realise that computing time is the bottleneck in AFM screens. Computing time dramatically increases with the length of the protein sequence. When using on the same GPU (NVIDIA GPU V100), for instance, it took 17 minutes to predict five models for a 183 amino acid (∼20k Da) protein pair, while it took 573 minutes to predict five models for a 1,726 amino acid (∼190 kDa) protein pair (**Figure 3**). The increase in prediction time by factor 34 for an increase in protein size by factor 9, illustrates the non-linear increase in computing times, which is also reported in the original AlphaFold2 manuscript (Jumper et al., 2021).

**Figure 3.**
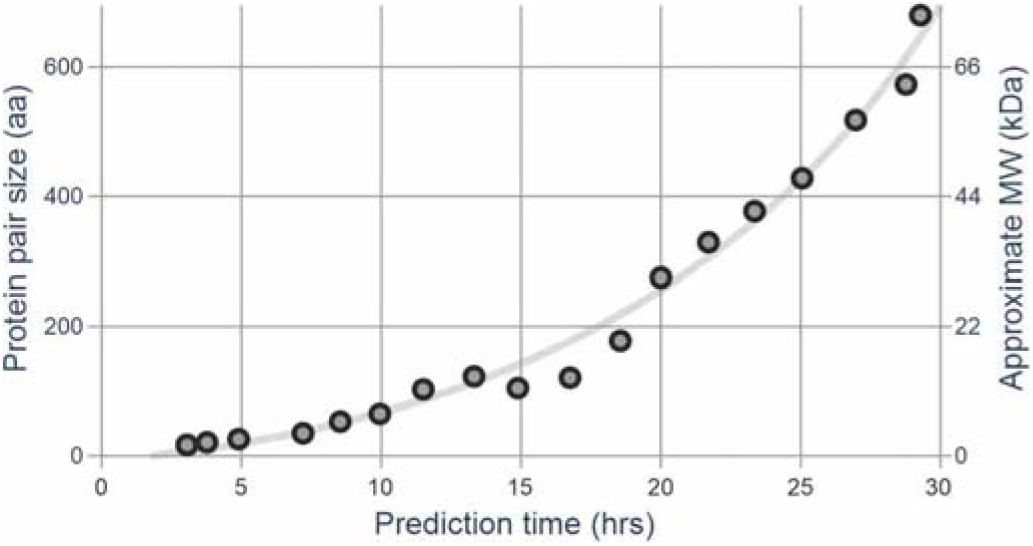
Small proteins model faster. Computing times on a single GPU (the NVIDIA GPU V100) were plotted for 18 random hydrolase-SSP protein pairs picked based on their different total residue length. The approximate molecular weight (MW) was calculated from residue length using 110 Da for each residue.

### Curate the input sequences

Since computing time is a bottleneck, considerable time will be saved by curating the input sequences to include only those that seem biologically relevant for the interaction, are not redundant to each other, or do not contain errors. For instance, our original screen of >11k protein pairs could have been substantially reduced if we had preselected for pathogen-derived proteins for which transcripts have been detected during infection. Likewise, sequences sharing e.g. >90% sequence identity can be removed to avoid the repeated prediction of highly similar complexes. Furthermore, non-natural residues (e.g. u) and the stop codon (*) should be removed since AFM can only handle the 20 natural amino acids.

### Remove irrelevant domains

Proteins have a modular structure and not all domains are present in the mature protein. Most hydrolases, for instance, have a signal peptide and a prodomain, which are both removed *in vivo* by protein processing. Removing irrelevant domains will reduce the computing times. In addition, these additional domains may obstruct the discovery of putative complexes. For instance, screens with prodomain-containing hydrolases may not result in the identification of inhibitors because the prodomain may occupy the substrate binding groove. For example, the complex between protease Pip1 and inhibitor EpiC2B is not predicted when the prodomain of Pip1 is included, since the prodomain of Pip1 occupies the substrate binding groove, obstructing EpiC2B binding (**Figure 4**). In other cases, however, other domains might be relevant for the interactions, so it is important to know which biologically relevant domains to include in the screen.

**Figure 4.**
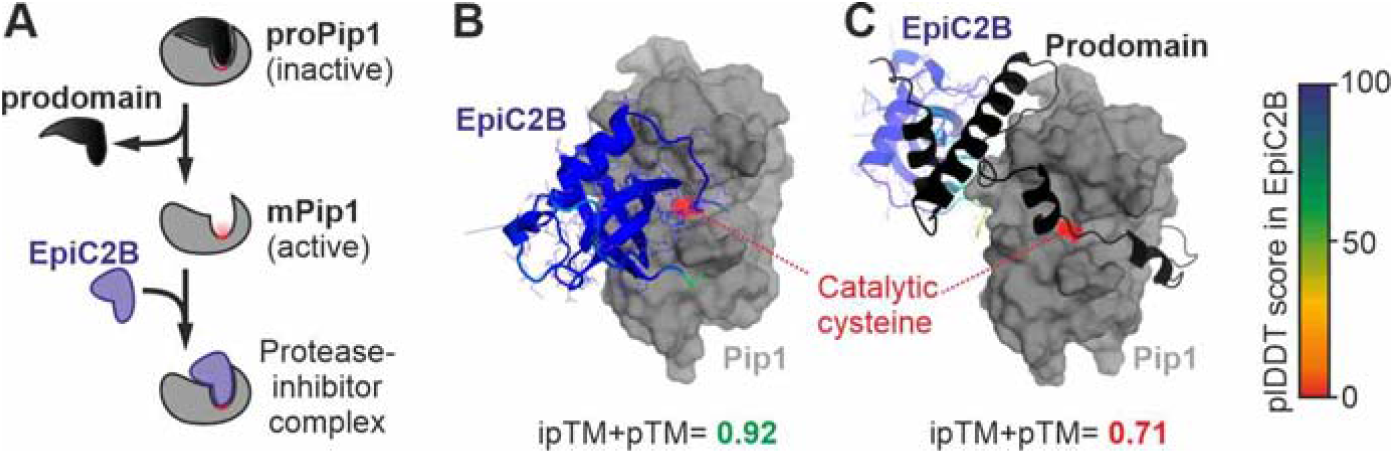
Prodomain prohibits inhibitor binding. **(A)** The N-terminal prodomain of Pip1 is proteolytically removed during Pip1 activation. **(B)** The complex predicted with AFM for EpiC2B with Pip1 shows the classic binding of a cystatin (EpiC2B) with papain (Pip1) (grey), associated with a predicted high score. **(C)** The complex predicted with AFM for EpiC2B with proPip1 shows that the substrate binding groove of Pip1 is occupied with the prodomain (black), displacing EpiC2B. The EpiC2B structure is shown as cartoon with the plDDT score in reverse rainbow colours; the mature protease of Pip1 is shown in grey as surface presentation with catalytic residue in red.

### Include positive controls

Positive controls of known interactors with the target protein are helpful to benchmark the output and determine the search parameters. For instance, the P69B-Epi1a and Pip1-EpiC2B complexes inspired us to look for similar SSPs that have an intrinsic fold and occupy the substrate binding groove of the proteases (**Figure 1**, step 5). Our screen also identified eight additional Kazal-like inhibitors for P69B and five additional cystatin-like inhibitors of Pip1. Although these 13 hits were retrospectively predictable from their sequence homology, identifying these inhibitors in our AFM assay independently increased the confidence in our screen.

***Include negative controls –***

Negative controls are important to identify an expected rate of false positives. The incompatible Epi1a-Pip1 and EpiC2B-P69B protein pairs, for instance, have ipTM+pTM scores of 0.48 and 0.47, respectively, indicating that scores <0.50 are below the cuttoff for reliable hydrolase-inhibitor pairs. Likewise, we included additional known SSP-hydrolase complexes in incompatible protein pairs to create a matrix with many negative controls that indicate the predicted scores of unlikely complexes (**Figure 5**). These datapoints can be used to set define the threshold in the screen. One caveat is of course that many of these negative controls are presumed but not proven and dual functionality of inhibitors is not uncommon (Grosse-Holz & Van der Hoorn, 2016).

**Figure 5.**
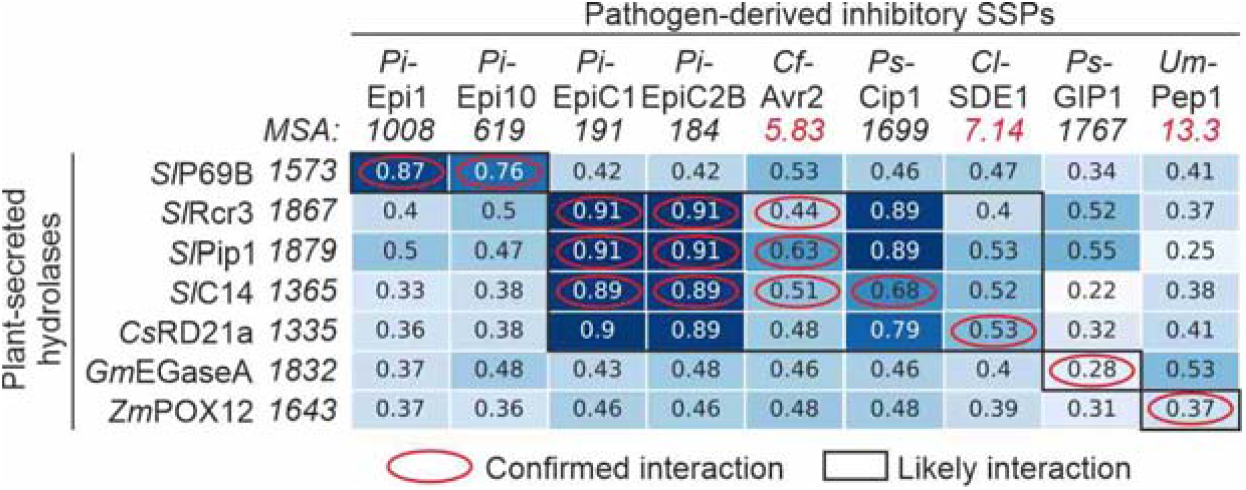
Using proteins from known complexes as negative and positive controls. Experimentally confirmed interactions between pathogen-secreted SSPs and plant-secreted hydrolases were analysed by AFM on the cluster in all possible SSP-hydrolase combinations to generate a heat map containing both positive controls (red circles) and negative controls (outside black boxes). The numbers in italics show the depth of the mean non-gap multiple sequence alignment (MSA) for each protein. Experimental evidence for SSP-hydrolase interactions have been published for *Pi*Epi1-*Sl*P69B (Tian et al., 2004), *Pi*Epi10-*Sl*P69B (Tian et al., 2005), *Pi*EpiC1/*Pi*EpiC2B-*Sl*Rcr3 (Song et al., 2009), EpiC1/EpiC2B-*Sl*Pip1 (Tian et al., 2007), *Pi*EpiC1/*Pi*EpiC2B-*Sl*C14 (Shabab et al., 2008); *Cf*Avr2-*Sl*Rcr3 (Rooney et al., 2003), *Cf*Avr2-*Sl*Pip1 (Shabab et al., 2008), *Cf*Avr2-*Sl*C14 (van Esse et al., 2008), PiCip1-Rcr3/Pip1/C14 (Shindo et al., 2016); *Cl*SDE1-CsRD21a (Clark et al., 2018), *Ps*GIP1-*Gm*EGaseA (Rose et al., 2002) and *Um*Pep1-*Zm*POX12 (Hemetsberger et al., 2012).

### Recycle Multiple Sequence Alignments (MSAs)

Every complex prediction involves generating a multiple sequence alignment (MSA) of both proteins before the complex is modelled. AFM would therefore normally generate two MSAs per protein pair, and would repeat this procedure for each new protein pair. The MSAs are, however, specific to the protein and not to the complex so reusing MSAs can significantly reduce computing times. Larger screens of e.g. 100 x 100 proteins (10,000 protein pairs) would therefore only require 200 MSAs (100 + 100), instead of 20,000 MSAs (10,000 x 2) that AFM would normally generate. Re-using MSAs can therefore reduce the MSA computing times by 99% in this example (**Figure 1**, step 3). This is possible by taking advantage of the modularity of AFM and use automated workflows to re-use MSAs for different complex predictions. Similar procedures have been implemented in AlphaPulldown (Yu et al., 2023). In addition, MSA generation can be quicker and still sufficiently accurate with MMSeqs2 (Steinegger & Söding, 2017), as used by ColabFold (Mirdita et al., 2022) instead of the initially used JackHMMER and HHblits in AFM (Evans et al., 2021).

### Control data storage

AFM generates many files for every protein pair, including a score file (.json), five models both before and after amber relaxation (.pdb), and metadata describing protein features and output statistics of the predictions (.pkl). Together, these files can amount to a gigabyte in storage, which quickly results in terrabytes of data, taking up all available storage. This problem can be overcome by incorporating automatic file compression steps in the workflow to free up the storage. Alternatively, automatically or routinely deleting less relevant files during the AFM screen could also help to manage storage, for example by only retaining files associated with the highest-scoring model or deleting files that score below a set threshold.

### Separate CPU-from GPU-intense steps

AFM uses both central processing units (CPUs) and graphics processing units (GPUs). A more efficient use of these resources is achieved if the multiple sequence alignments (MSAs), which are run on CPUs, is separated from the 3D model prediction, which is run on GPUs. This way, only hardware required for the specific step can be requested. A similar separation of tasks over CPUs and GPUs has been implemented in AlphaPulldown (Yu et al., 2023). In our workflow, we reduced computing time considerably by using the workflow manager Snakemake (Köster & Rahmann, 2012) to compartmentalise the workflow into CPU- and GPU-specific steps.

### Try to get MSA >100

AFM uses Multiple Sequence Alignments (MSAs) to predict the protein fold (Jumper et al., 2021, **Figure 1**). Consequently, the accuracy of structural predictions increases with the number of available sequences. Confidence in structural predictions is usually high when MSA>100, but is generally poor when MSA<10. ColabFold (Mirdita et al., 2022) is efficient because it uses MMSeqs2 (Steinegger & Söding, 2017) to quickly generate MSAs from a reduced, clustered sequence database (ColabFoldDB, 738M entries). This is usually not a problem for plant proteins, but e.g. fungal and oomycete proteins that are not present in every sequenced organism, such as effectors, tend to have lower MSAs and are therefore poorly modelled using reduced databases. Running a custom AFM workflow on a computing cluster enables the use of a larger sequence database and the inclusion of additional sequences that may not be publicly available. This is illustrated by the increased coverage of pathogen-derived proteins with MSA<100 when adding a custom fungal database (35M entries) or including to the full ‘Big Fantastic Database’ (BFD, 2,500M entries, Jumper et al., 2021) (**Figure 6A**). Consequently, the modelling confidence increases significantly, demonstrated with increased plDDT scores (**Figure 6B**).

**Figure 6.**
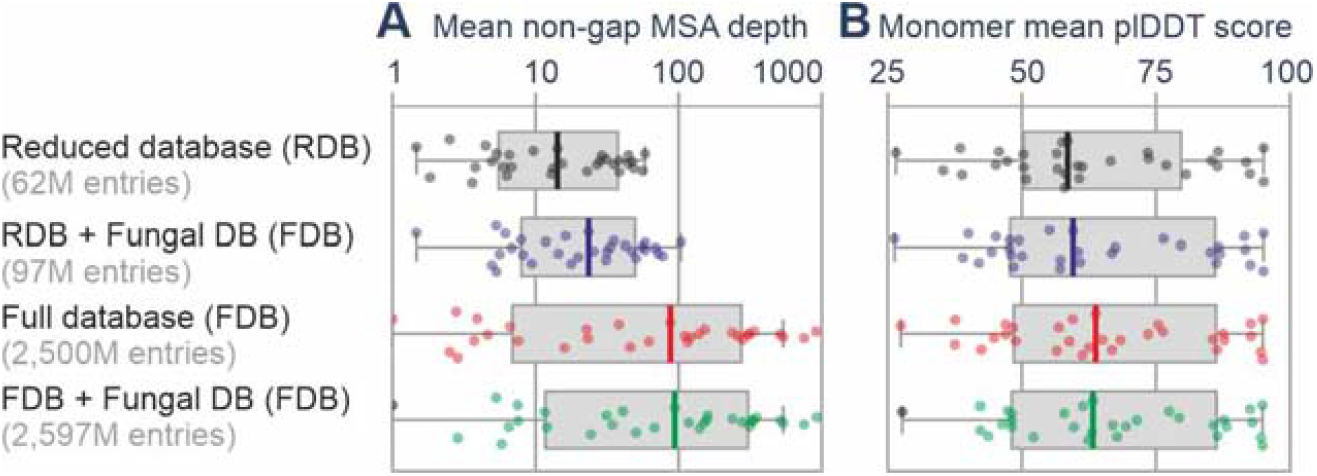
Expanding the sequence database increases modelling confidence. 32 fungal and oomycete proteins initially having a mean non-gap MSA depth<100 in the reduced database (RDB, 62M entries) were reanalysed with AFM upon extending the database with a fungal database (FDB, 35M entries), the full database (BFD, 2,500M entries), or both. **(A)** Mean non-gap MSA depth increases when expanding the database. **(B)** The modelling confidence, expressed as the mean plDDT score, increases with increased MSA.

### Evaluate the predicted scores

By default, AFM predicts five complexes for a given protein pair, each with their own scores. The produced models (.pdb files) can be visualised in ChimeraX or PyMol. The ranking is based on the combined predicted template modelling (pTM) and interface pTM (ipTM) scores, which are combined in a single score for each complex (0.2*pTM+0.8*ipTM), ranging from 0 (low) to 1 (high confidence). You can find this score in the .json file and can also be visualised via PAE Viewer (Elfmann & Stülke, 2023). AFM also infers the predicted local distance difference test (plDDT) score for every residue in the model. The plDDT score is usually visualised in structures with the reverse rainbow colour scheme ranging from red (low confidence) to blue (high confidence). The plDDT confidence scores are documented in the .pdb files and can be visualised in PyMol with command [`spectrum b, rainbow_rev, minimum=0, maximum=100`]. The mean non-gap Multiple Sequence Alignment (MSA) depth can be calculated with a script from the features.pkl file. ColabFold summarises the MSA depth in a graphic presentation but does not provide the .pkl file. Finally, the Predicted aligned error (PAE) gives a distance error for every pair of residues with values between 0-36 Å and can be analysed in PAE viewer (Elfmann & Stülke, 2023).

### Use ipTM+pTM to select candidate interactions

AFM calculates the predicted template modelling (pTM) and interface pTM (ipTM) for each predicted complex, and these are combined in a single score with 4-fold more weight on the ipTM score when compared to the pTM score (combined score = 0.8 ipTM + 0.2 pTM). We found that selecting scores above 0.75 results in good candidate complexes in which both proteins have a high-confidence predicted fold. For SSP-hydrolase protein pairs, a threshold of ipTM+pTM ≥ 0.75 resulted in the discovery of several novel protein-protein interactions (Homma et al., 2023).

### Beware of typical AFM errors

Rather than predicting separate models of two proteins and then docking them together, AFM predicts protein complexes by folding both proteins simultaneously. The advantage is that this may result in protein conformations that are different from when these proteins were folded separately. Occasionally, however, AFM predicts intertwined folds that are unlikely to occur in nature, for instance when part of one protein becomes an integral structural part of the other protein, e.g. by participating in a β-sheet structure. These instances are, however, quite rare (<5%). We also noticed that ColabFold online by default runs AFM without amber relaxation (Hornak et al., 2006) as final subroutine. This saves computing time but can generate models in which atoms occupy the same space.

### Beware of false negatives

We discovered that some experimentally confirmed hydrolase-inhibitor interactions did not yield high-confidence complex from AFM predictions. The fungal Avr2 protein, for instance, is known to inhibit Pip1 (Shabab et al., 2008), yet AFM predicts models that score only (0.2pTM+0.8ipTM=) 0.44 (**Figure 7**). This might be caused by the low sequence coverage of Avr2 (mean non-gap MSA depth = 5.82), hence AFM has limited power to infer co-evolving residues within Avr2. Interestingly, the AFM-predicted models also fail to adequately connect the disulphide bridges within Avr2, which have been determined experimentally (van ‘t Klooster et al., 2011). Feeding AFM with additional Avr2 sequences will presumably increase the sequence depth and improve the predictions of the Avr2-Pip1 complex. In addition, AFM also predicts low scores for *Cl*SDE1-RD21 (0.53); *Um*Pit2-CP1A (0.35) and *Um*Pep1-POX12 (0.37), which have all been experimentally shown to interact (Clark et al., 2018; Mueller et al., 2013; Hemetsberger et al., 2012). As with Avr2, the mean non-gap MSA depth is below 100 for these three effectors, indicating that sequence availability might be a common limit on their structural predictions.

**Figure 7.**
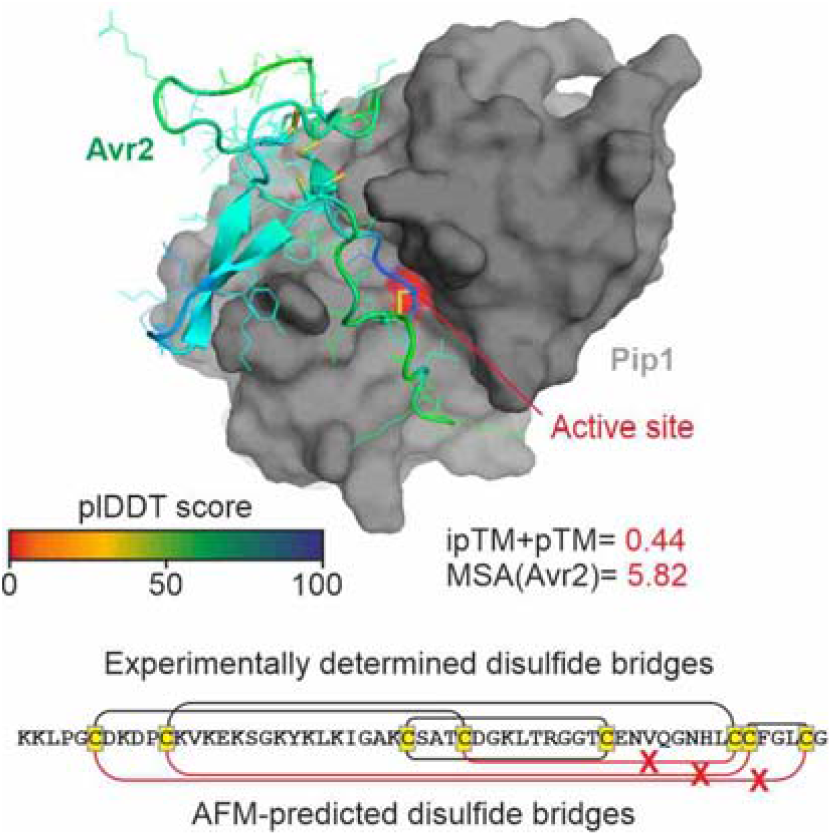
AFM predicts false negative Avr2-Pip1 complex. The experimentally validated complex of Avr2 and Pip1 (Shabab et al., 2008) was predicted with AFM on the computing cluster. The best model has a low ipTM+pTM score and has mis-predicted (red ‘X’) three of the four disulphide bridges that were experimentally determined (Van ‘t Klooster et al., 2011).

### Beware of false positives

Not all hydrolase-SSP interactions with ipTM+pTM ≥ 0.75 might be true inhibitors, even if the SSPs have an intrinsic fold that blocks access to the active site. To illustrate this, we tested the different Kazal domains of Epi1 and Epi10. Epi1 contains two Kazal domains of which the first (Epi1a) can inhibit P69B, but the second (Epi1b) cannot (Tian & Kamoun, 2005). Interestingly, highest scores for the Epi1a-P69B and Epi1b-P69B complexes are both above the 0.75 threshold, but the mean score for Epi1b is significantly lower (**Figure 8A**). Likewise, Epi10 contains three Kazal domains but only the second (Epi10b) is thought to inhibit P69B (Tian et al., 2005). Although the AFM score of Epi10b is the highest, AFM predictions are also high for Epi10a and Epi10c (**Figure 8B**). We also tested engineered Pip1 (ePip), which we recently developed by substituting two residues (D147R, V150E) to avoid its interaction with EpiC2B (Schuster et al., 2023). Importantly, AFM is unable to distinguish Pip1-EpiC2B from ePip-EpiC2B (**Figure 8C**). This is consistent with the notion that AFM is unable to predict the effect of non-natural mutations (Buel & Walters, 2022). Thus, AFM can produce false positive hits that cannot be confirmed experimentally. Nevertheless, these false positives indicate that homologous proteins and/or mutants might interact experimentally.

**Figure 8.**
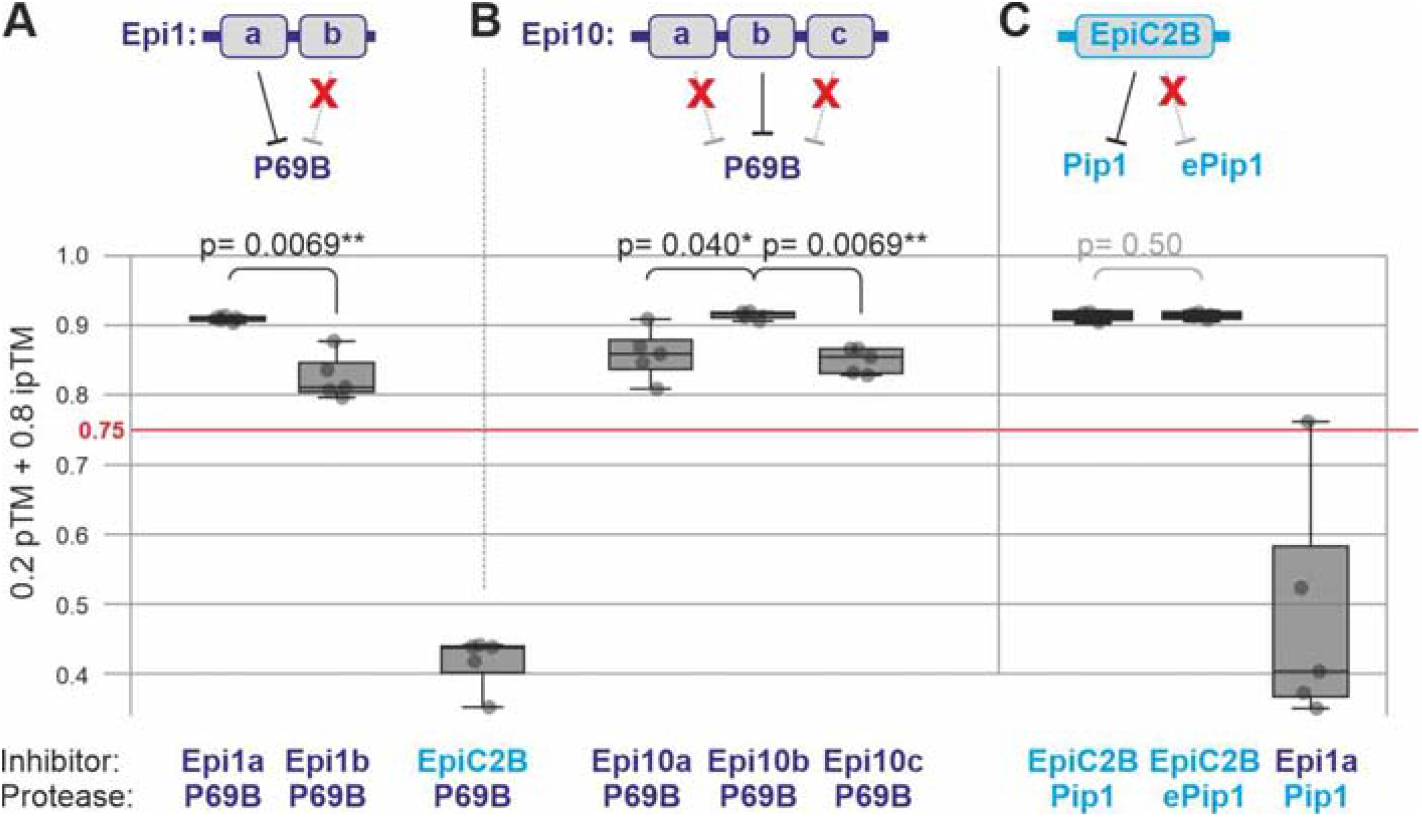
AFM predicts false positive protease-inhibitor interactions. **(A)** Of the two Kazal-like domains in Epi1, only the first (Epi1a) inhibits P69B, whilst the second (Epi1b) does not inhibit P69B (Tian & Kamoun, 2004). However, AFM predicts complexes with scores above the 0.75 threshold, yet the false positive scores are lower than the true positive scores.**(B)** Of the three Kazal-like domains in Epi10, only the middle (Epi10b) is thought to inhibit P69B, whilst the others do not inhibit P69B (Tian et al., 2005). However, AFM predicts complexes with scores above the 0.75 threshold, yet the false positive scores are lower than the true positive scores.**(C)** Engineered Pip1 (ePip1) carries two substitutions that abolish inhibition by EpiC2B (Schuster et al., 2024). Yet, the scores predicted for these two complexes are both above the 0.75 threshold and are statistically indistinguishable.

### Explore hits manually

We analysed complexes with ipTM+pTM ≥ 0.75 manually in PyMol. This process was accelerated by using a script that generated PyMol (.pse) files that highlighted the active site of the hydrolase so that the relative position of the SSP on the hydrolase could be quickly determined. Manual evaluation by two independent researchers of a few hundred complexes takes only a few days and produced consistent observations. Manual evaluation is a useful exercise as we were able to identify complex features and use several important parameters, such as the confidence of the SSP fold and how well it is predicted to interacts with its target. Loose associations of the SSP with the hydrolase and the absence of disulphide bridges in Cys-rich SSPs are suspicious for being false positives since many SSPs have disulphide bridges to stabilise them in the apoplast. We disregarded most complexes as inhibitor candidates because they did not block the active site or lacked an intrinsic structure. Only 10% of the high scoring complexes seemed candidate inhibitors that interacted in a similar way to known hydrolase inhibitors. We have tested automation of this evaluation by determining if the active site is in the interface and how the plDDT and PAE scores are at the interface but were unable to establish robust parameters that would have led to the confirmed inhibitors that were discovered manually.

### Remaining challenges

Structural predictions by AI are quickly improving. At this stage, AFM is unable to predict the effect of single site substitutions (Buel & Walters, 2022). AFM is also not yet able to consider the impact of a different pH, presence of ions and oxidative burst on proteins. Likewise, the impact of post-translational modifications (PTMs) e.g. by phosphorylation, methylation or glycosylation, are important parameters to include in predictions. AFM is also unable to predict interactions with small molecules and nucleic acids, although this area is rapidly developing. Another unresolved issue is the pairing of co-evolving proteins from different species, which could be used to strengthen the predictions of protein complexes. Weaker, temporal interactions involving small interfaces may require the selection of different parameters than ipTM+pTM scores, such as the Local Interaction Scores (LOS) extracted from the Predicted Alignment Error (PAE, Kim et al., 2023). We also need standards in this new field to calculate and compare the strengths of predicted protein-protein interactions and include the prediction files as supplemental information for publications. Ultimately, we should be able to predict targets for agrochemicals and pathogen-derived toxins and use AI to understand the dynamics of interkingdom interactions of plants with symbionts and pathogens. Excitement looms ahead.

### AlphaFold 3 (!)

After the completion of this manuscript, AlphFold 3 was released (Abramson et al., 2024). AlphaFold 3 offers additional features such as interactions with nucleic acids and small molecules, as well as post-translational modifications (PTMs). AlphaFold 3 is also faster than AlphaFold2. However, we found that ipTM+pTM scores for complexes of *Sl*P69B with *Cf*Ecp36, *Fo*Six15 and *Fo*TIL predicted by AlphaFold 3 are still below the 0.75 threshold, in contrast to our customised AFM workflow. We also noticed that AlphaFold 3 still does not correctly predict the disulphides in Avr2 in the Avr2-Rcr3 complex. In addition, AlphaFold 3 only runs on Google servers with a maximum 20 tasks per day and source code has not yet been made available to implement local AlphaFold 3 workflows However, this area is developing quickly and solutions will certainly be found.

## Acknowledgements

We like to thank Sophien Kamoun for useful suggestions and the Advanced Research Computing (ARC, Richards, 2015) facility of the University of Oxford for access to their high-performance computing cluster. This project was financially supported by the Interdisciplinary Doctoral Training Program (DTP) of the BBSRC (project DDT00230, JL), and the European Research Council ERC-AdG-2020 grant ‘ExtraImmune’ (project 101019324, FH, RH).

